# Differential gene expression in *Drosophila melanogaster* and *D. nigrosparsa* infected with the same *Wolbachia* strain

**DOI:** 10.1101/2020.02.19.941534

**Authors:** Matsapume Detcharoen, Martin P. Schilling, Wolfgang Arthofer, Birgit C. Schlick-Steiner, Florian M. Steiner

**Affiliations:** Molecular Ecology Group, Department of Ecology, University of Innsbruck

**Keywords:** endosymbionts, host-microbe interactions, RNA sequencing, symbiosis, transcriptomics

## Abstract

*Wolbachia*, maternally inherited endosymbionts, infect nearly half of all arthropod species. *Wolbachia* manipulate their hosts to maximize their transmission, but they can also provide benefits such as nutrients and resistance to viruses for their hosts. The *Wolbachia* strain *w*Mel was recently found to increase locomotor activities and possibly trigger cytoplasmic incompatibility in the fly *Drosophila nigrosparsa*. Here, we compared differential gene expression in *Drosophila melanogaster* (original host) and *D. nigrosparsa* (novel host), both uninfected and infected with *w*Mel, using RNA sequencing to see if the two Drosophila species respond to the infection in the same or different ways. A total of 2164 orthologous genes were used. We found species-specific gene expression patterns. Significant changes shared by the fly species were confined to the expression of genes involved in heme binding and oxidation-reduction; the two host species differently changed the expression of genes when infected. Some of the genes were down-regulated in the infected *D. nigrosparsa*, which might indicate small positive effects of *Wolbachia*. We discuss our findings also in the light of how *Wolbachia* survive within both the native and the novel host.

## Introduction

Change in gene expression underlies various essential phenomena in organisms. Different gene expression patterns reveal how organisms respond to different environments. Tools such as quantitative PCR, microarrays, and high-throughput RNA sequencing have been developed to detect differences in gene expression. RNA sequencing is a powerful method to study the transcriptomic function of organisms such as differential gene expression because of its wide range of applications (Stark *et al*., 2019). Recently, differential gene expression using RNA sequencing has been widely used to study relationships between hosts and their endosymbionts (e.g. Gutzwiller *et al*., 2015; Bennuru *et al*., 2016; Baião *et al*., 2019).

*Wolbachia* (Alphaproteobacteria) are a group of bacterial endosymbionts found in arthropods and nematodes (Werren *et al*., 2008). It is estimated that around half of all arthropods are infected with *Wolbachia* (Zug and Hammerstein, 2012; Sazama *et al*., 2017), with their strain diversity estimated at around 100,000 strains (Detcharoen *et al*., 2019). By maternal transmission, these endosymbionts manipulate their hosts for their benefits in various ways, such as feminization, cytoplasmic incompatibility, male-killing, and parthenogenesis (Werren *et al*., 2008). In some cases, *Wolbachia* also provide benefits to their hosts, such as when supplying vitamins to *Cimex lectularius* bedbugs (Hosokawa *et al*., 2010) and virus protections in *Drosophila* species (Hedges *et al*., 2008; Teixeira *et al*., 2008; Osborne *et al*., 2009; Cattel *et al*., 2016).

There are around 2000 *Drosophila* species (O’Grady and DeSalle, 2018) ranging from habitat generalists to habitat specialists (Kellermann *et al*., 2009). The model organism *Drosophila (Sophophora) melanogaster* is a generalist with a cosmopolitan distribution (Bächli *et al*., 2004). *Drosophila (Drosophila) nigrosparsa* is an alpine species found at around 2000 m above sea level in Central and Western Europe (Bächli *et al*., 2004). Due to the habitat specificity of *D. nigrosparsa*, molecular and physiological traits and potential effects of warming temperatures on this species have been studied (Arthofer *et al*., 2015; Kinzner *et al*., 2016, 2018, 2019; Cicconardi *et al*., 2017; Tratter Kinzner *et al*., 2019). Wild populations of *D. melanogaster* are commonly infected with *Wolbachia* (Verspoor and Haddrill, 2011), while no wild population of *D. nigrosparsa* infected with *Wolbachia* has been found to date (M. Detcharoen, unpubl.). We recently transinfected the *Wolbachia* strain *w*Mel from *D. melanogaster* into *D. nigrosparsa* and studied several traits including *Wolbachia* density as well as host temperature tolerance, larval and adult locomotion, and cytoplasmic incompatibility (Detcharoen *et al*., 2020). Our analysis of the new host-endosymbiont system of *Drosophila nigrosparsa* infected with *Wolbachia w*Mel revealed increased locomotion compared with flies tetracycline cured from their infection as well as hints of weak cytoplasmic incompatibility (Detcharoen *et al*., 2020).

In searching for molecular mechanisms behind the increased locomotion and cytoplasmic incompatibility in *D. nigrosparsa*, we here used differential gene expression analysis using RNA sequencing. We aimed to investigate the effects of *Wolbachia w*Mel on gene expression in *D. melanogaster*, the native host, and *D. nigrosparsa*, the novel host, for a comparative analysis to find out whether these two species respond to the infection in the same or in different ways.

## Experimental Procedures

### Fly culture

Uninfected *Drosophila melanogaster* (mu_0) were provided by Luis Teixeira as well as *D. melanogaster* infected with *Wolbachia* strain *w*Mel. From the latter, three infected lines, that is, mi_1, mi_2, and mi_3, were generated. The uninfected *D. melanogaster* line was kept in the laboratory for about six generations and 30 generations for the infected lines prior to the experiment.

As uninfected *D. nigrosparsa* (nu_0), individuals of isofemale line iso12 were used that had been established using a population at Kaserstattalm, Tyrol, Austria (47.13°N, 11.30°E) in 2010 (Arthofer *et al*., 2015; Cicconardi *et al*., 2017) and had been maintained in the laboratory for about 60 generations before the founding of line nu_0. *w*Mel-infection of *D. nigrosparsa* was achieved as described in (Detcharoen *et al*., 2020). Briefly, cytoplasm of *D. melanogaster* containing *Wolbachia w*Mel was transinfected into embryos of *D. nigrosparsa* line nu_0, and three infected lines, ni_3, ni_6, and ni_8, were generated.

Both fly species were cultured as described in Kinzner *et al*. (2018). Briefly, *Drosophila melanogaster* was cultured using corn food at a density of 80 embryos per vial with 8 ml food. For *D. nigrosparsa*, approximately 50 males and 50 females were put in a mating cage supplied with grape-juice agar, malt food, and live yeast. Embryos and larvae were collected and put in glass vials with 8 ml malt food at a density of 80 embryos per vial. All fly stocks were maintained at 19 °C, 16 h: 8 h light: dark cycle, and 70% relative humidity in an incubator (MLR-352H-PE, Panasonic, Japan).

Five females per line were randomly collected prior to the experiment to check for *Wolbachia* infection. DNA was extracted, and the infection of each line in the generation used for RNA analyses was confirmed using PCR with the primers wsp81F and wsp691R (Braig *et al*., 1998). Besides the infection of lines in both species with *Wolbachia*, all fly lines used in this study were found not to be infected with other known bacterial endosymbionts (data not shown).

### RNA extraction and sequencing

Because of the relatively long development time of *D. nigrosparsa*, fourteen-day old females of *D. nigrosparsa* and five-day old females of *D. melanogaster* of both uninfected and infected lines (totaling eight lines) were randomly collected using short carbon-dioxide anesthesia. Only female flies were used because *Wolbachia* are maternally transmitted. All flies were killed by snap freezing in liquid nitrogen within 5 minutes after taking them from their regular regimes as described in the previous section and at the same time of the day (at 1100 hours a.m.) to control for any circadian-rhythm based variation in gene expression. RNA from individual flies was extracted using RNeasy Micro Kit (Qiagen, Hilden, Germany) following the manufacturer’s protocol. Quantity of RNA was measured using Quant-iT RiboGreen RNA Assay Kit (Thermo Fisher Scientific, USA). RNA extracts of five individuals belonging to the same line were pooled to have a minimum RNA content of 2.5 µg per replicate and five replicates per fly line. RNA library preparations and sequencing using Illumina NextSeq 500 were done at IGA Genomics (Udine, Italy).

### Sequence alignment and differential expression analyses

Single-end raw reads (SRA database, BioProject PRJNA602188, accession number SAMN13885146-SAMN13885185) were subjected to quality-check with FastQC version 0.11.8 (Andrews, 2010). Trimmomatic version 0.38 (Bolger *et al*., 2014) was used to remove adapters including the first and the last three nucleotides, to cut reads when the average Phred score dropped below 20 in a 4-bases sliding window, and to drop reads shorter than 40 bases. Reads were quantified relative to their reference transcriptomes using Salmon version 0.12.0 (Patro *et al*., 2017), that is, reads belonging to *D. melanogaster* were mapped to the reference transcriptome of *D. melanogaster* build BDGP6, and reads belonging to *D. nigrosparsa* were mapped to its previously published transcriptome (Arthofer *et al*., 2015). The numbers of quantified reads were imported to R (R Core Team, 2018) using the package tximport (Soneson *et al*., 2016).

The mapped genes of *D. nigrosparsa* were translated to protein sequences and blasted to *D. melanogaster* using tblastn function implemented in Flybase (Thurmond *et al*., 2019) to search for orthologous genes. Only orthologous genes shared between the two species were used for further analyses. Genes with less than 50 counts across all samples were removed. BaySeq (Hardcastle and Kelly, 2010) was used to analyze differential expression between uninfected and infected flies of each species. Priors were estimated using a negative binomial distribution with quasi-maximum-likelihood, and posterior likelihoods for each orthologous gene were calculated. Genes were sorted by their posterior likelihood, and those in the third quartile were selected. Gene ontology (GO) analyses were done using DAVID version 6.8 (Huang *et al*., 2009), and the Benjamini-Hochberg procedure was used to control for false-discovery rate using an alpha value of 0.05. Normalized read counts of genes with posterior probabilities greater than 0.5 were grouped based on Pearson correlation and visualized with heatmaps generated using the R package NMF (Gaujoux and Seoighe, 2010). Principal component analysis was performed based on read counts, and relationships among samples were visualized with biplots. The R package vegan (version 2.5-4) (Oksanen *et al*., 2019) was used to calculate analysis of similarities (ANOSIM) among samples based on infection status, lines, and species and non-metric multidimensional scaling (NMDS) to visualize the similarities. All analyses were done in R (R Core Team, 2018), and visualizations were created using the package ggplot2 (Wickham, 2016).

## Results

After quality control, an average of (mean ± standard deviation) 24.8 ± 7.1 and 23.4 ± 4.9 million high-quality reads were found per replicate (pooling five individuals) of *D. melanogaster* and *D. nigrosparsa*, respectively. About 85% of the *D. melanogaster* reads and 80% of the *D. nigrosparsa* reads were mapped to their respective reference transcriptome. We removed the uninfected *D. melanogaster* replicate sample mu_0.1 from the analyses because of a contamination of male flies. A total of 2164 genes in *D. nigrosparsa* were found to be orthologous in *D. melanogaster*. After removing genes with low expression across samples, 2084 genes remained in the dataset.

### Variation among pools of individuals within replicate lines and among replicate lines within species

Following differential expression analysis, genes were ordered according to their posterior probabilities (Figure S1). We found 304 and 367 genes with posterior probabilities in the third quartile in *D. melanogaster* and *D. nigrosparsa*, respectively (Table S1). Among these, 87 genes were shared between the two species (Figure 1A). We found 103 and 87 genes with a posterior probability > 0.5 in *D. melanogaster* and in *D. nigrosparsa*, respectively.

**Figure 1.**
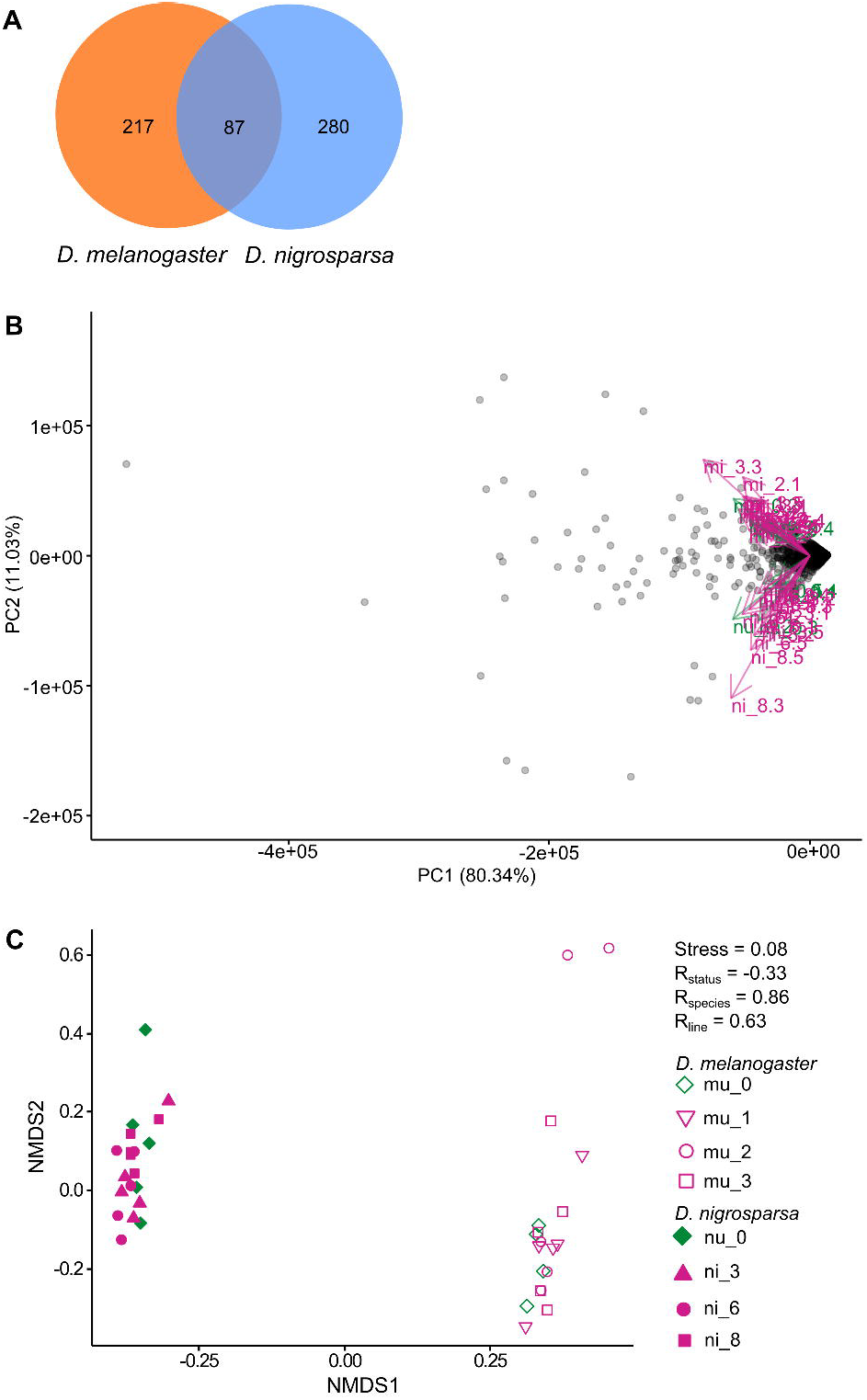

We found strong variation among pools of individuals but not between infection status when compared within and between species, such as in infected-*D. melanogaster* sample mi_3.3 and in uninfected-*D. nigrosparsa* sample nu_0.2. These samples appeared to have more highly expressed genes than the remaining samples (Figure 2). Within species, we observed more variation among lines (ANOSIM, *D. melanogaster* R = 0.14, *D. nigrosparsa* R = 0.33) than between infection status (ANOSIM, *D. melanogaster* R < 0.01, *D. nigrosparsa* R = 0.04) (Figure S2).

**Figure 2.**
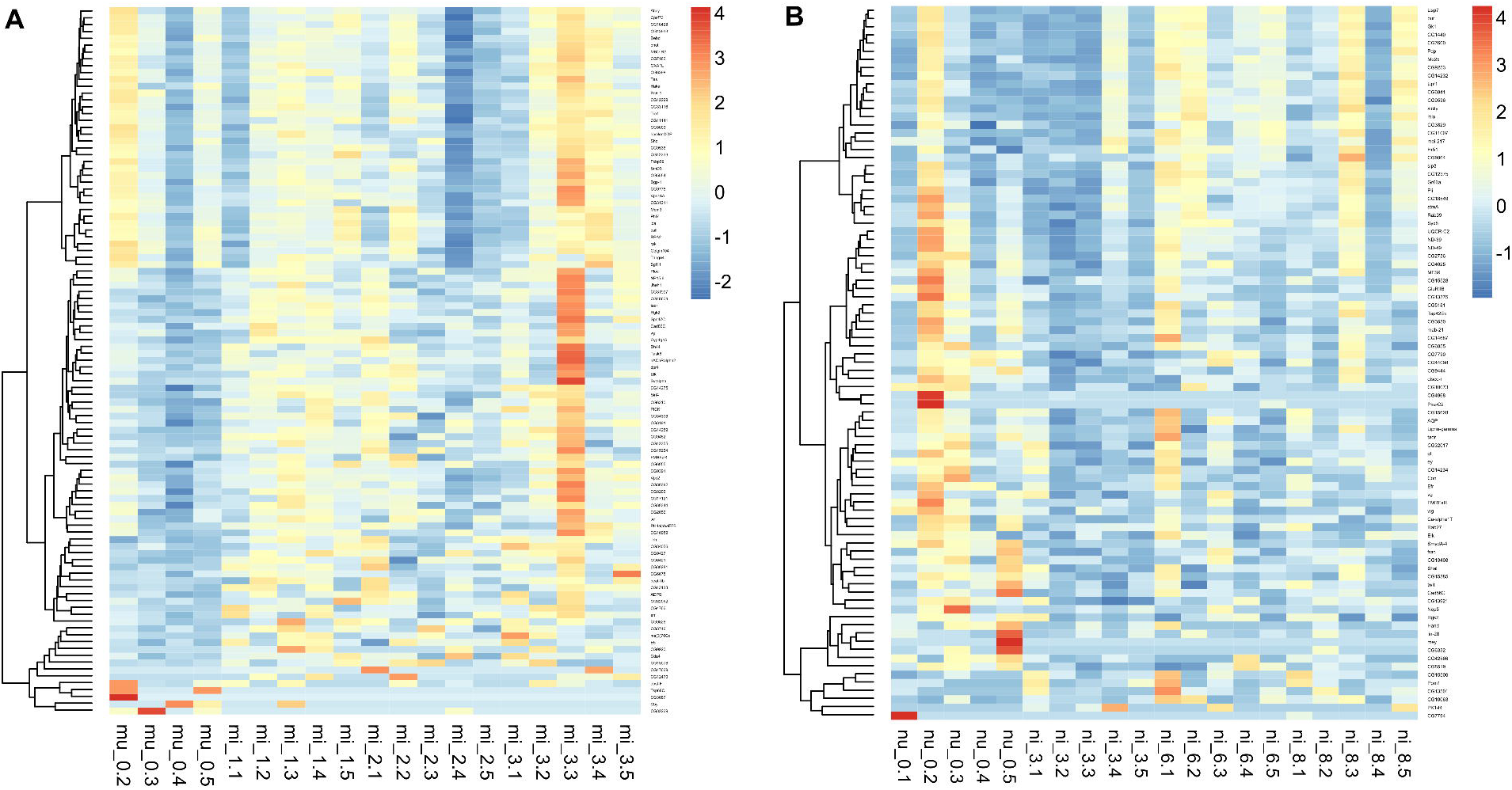

### Strong difference between species

Gene expression in *D. melanogaster* strongly differed from that in *D. nigrosparsa* (Figure 1B). In the PCA, PC1 explained most of the variation among orthologous genes in both species (80.34%). As supported by the ANOSIM results (Figure 1C), we observed a clear separation between species regarding the infection status (R = 0.86).

### Functional analysis of orthologous genes

We used orthologous genes of both species for GO term analysis. Fifteen GO terms were significantly enriched in *D. melanogaster* and sixteen terms in *D. nigrosparsa* (Table 1). Among these, two GO terms, heme binding (molecular function) and oxidation-reduction process (biological process), were shared between the two species. Several genes belonging to these two shared GO terms encode proteins that belong to cytochrome P450 families (Table 2). Genes belonging to the molecular function heme binding were likely to be expressed more highly in *Wolbachia*-infected than uninfected *D. melanogaster*, but there was no such trend in *D. nigrosparsa* (Table 2). Two heme-binding genes (*snl* and *Cyt-c1L*) were shared between the two species, and these were likely to be expressed more highly in infected than in uninfected flies.

**Table 1.**
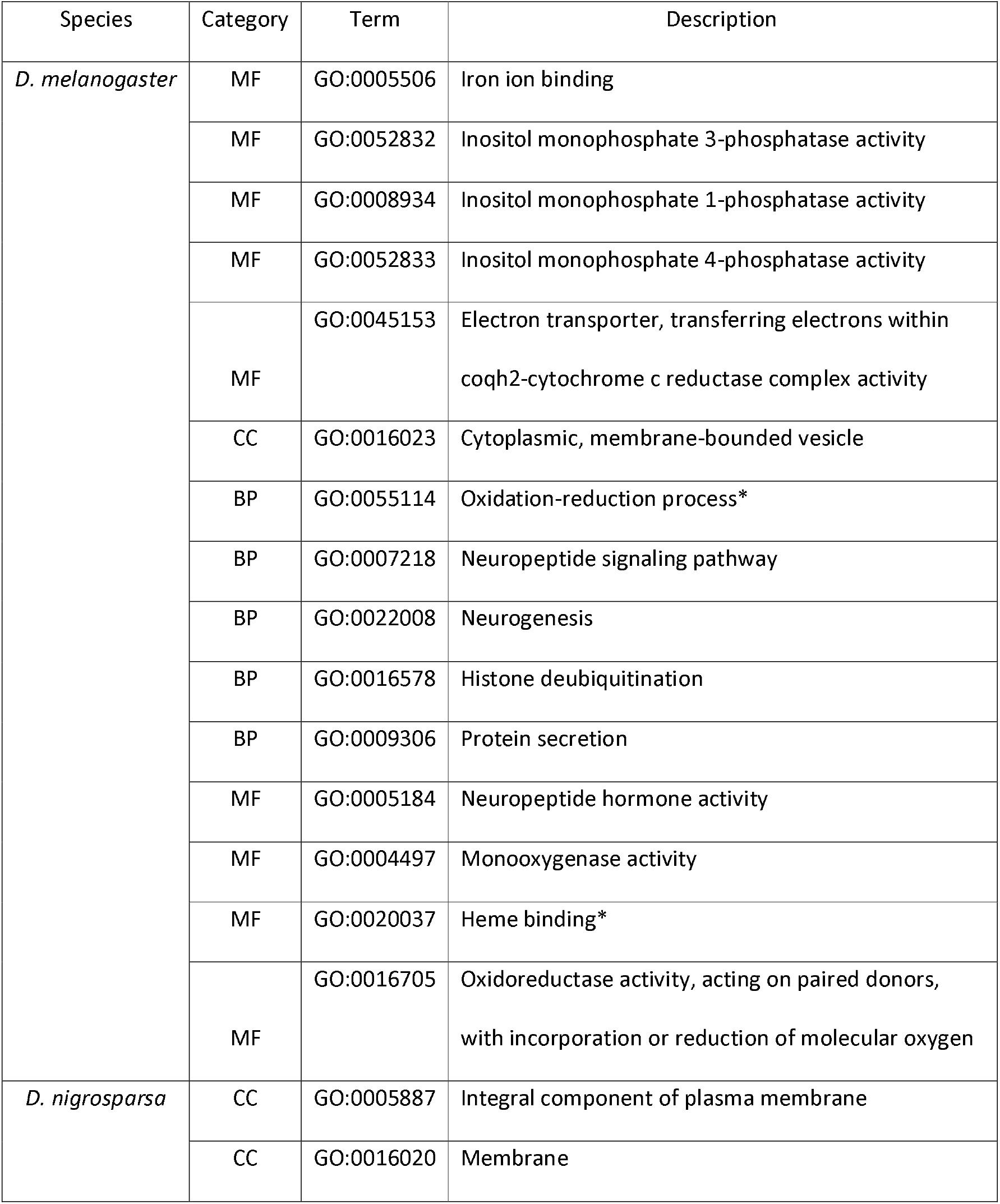
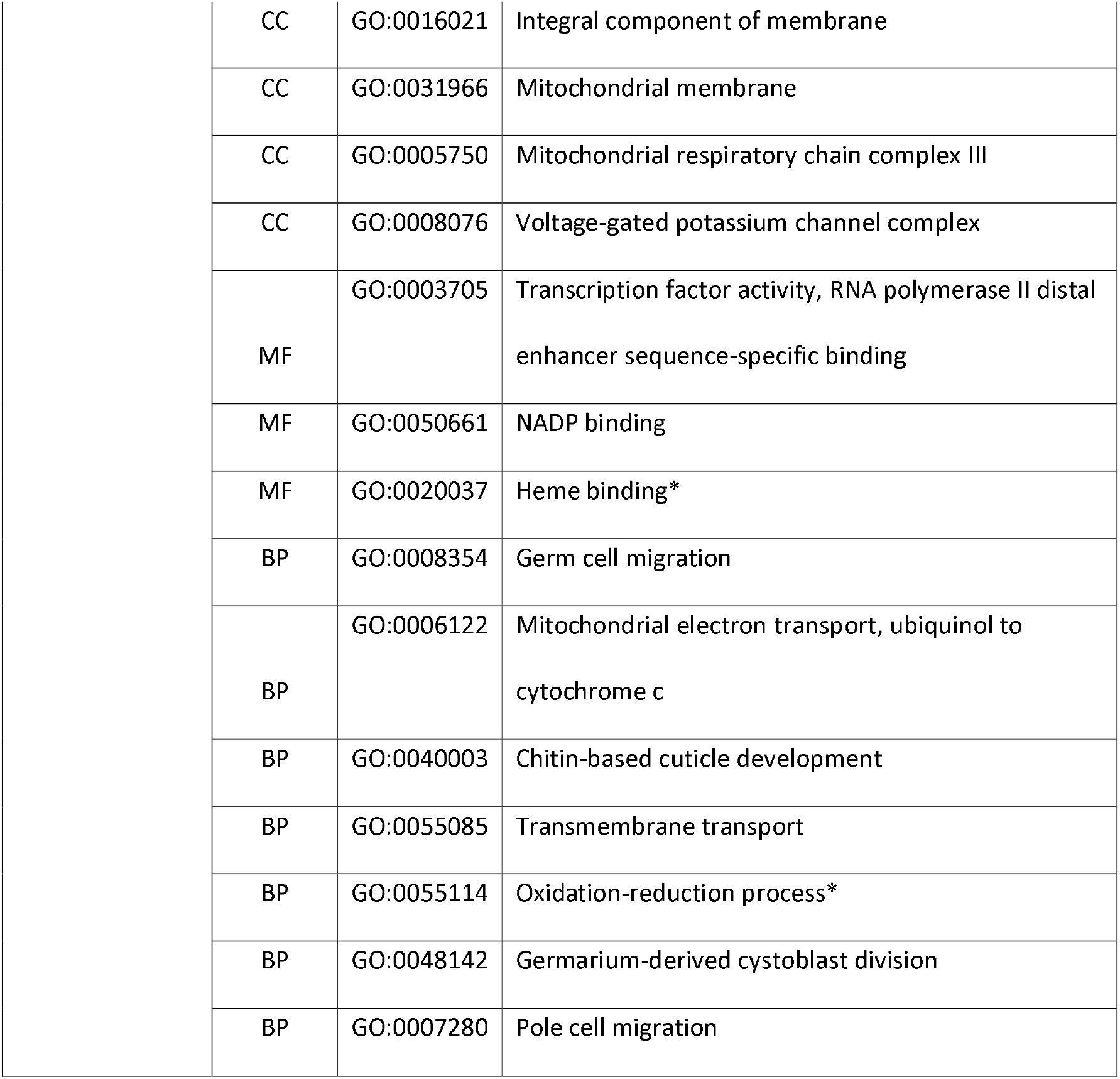
Significant gene ontology (GO) terms in *Drosophila melanogaster* and in *D. nigrosparsa* using all orthologous genes. Two GO terms were shared between the two species, heme binding and oxidation-reduction process (marked with ^*^). MF, CC and BP are molecular function, Cellular component, and molecular function, respectively.

| Species | Category | Term | Description |
| --- | --- | --- | --- |
| *D. melanogaster* | MF | GO:0005506 | Iron ion binding |
|  | MF | GO:0052832 | Inositol monophosphate 3-phosphatase activity |
|  | MF | GO:0008934 | Inositol monophosphate 1-phosphatase activity |
|  | MF | GO:0052833 | Inositol monophosphate 4-phosphatase activity |
|  | MF | GO:0045153 | Electron transporter, transferring electrons within coqh2-cytochrome c reductase complex activity |
|  | CC | GO:0016023 | Cytoplasmic, membrane-bounded vesicle |
|  | BP | GO:0055114 | Oxidation-reduction process* |
|  | BP | GO:0007218 | Neuropeptide signaling pathway |
|  | BP | GO:0022008 | Neurogenesis |
|  | BP | GO:0016578 | Histone deubiquitination |
|  | BP | GO:0009306 | Protein secretion |
|  | MF | GO:0005184 | Neuropeptide hormone activity |
|  | MF | GO:0004497 | Monooxygenase activity |
|  | MF | GO:0020037 | Heme binding* |
|  | MF | GO:0016705 | Oxidoreductase activity, acting on paired donors, with incorporation or reduction of molecular oxygen |
| *D. nigrosparsa* | CC | GO:0005887 | Integral component of plasma membrane |
|  | CC | GO:0016020 | Membrane |

|  |  |  |
| --- | --- | --- |
| CC | GO:0016021 | Integral component of membrane |
| CC | GO:0031966 | Mitochondrial membrane |
| CC | GO:0005750 | Mitochondrial respiratory chain complex III |
| CC | GO:0008076 | Voltage-gated potassium channel complex |
| MF | GO:0003705 | Transcription factor activity, RNA polymerase II distal enhancer sequence-specific binding |
| MF | GO:0050661 | NADP binding |
| MF | GO:0020037 | Heme binding* |
| BP | GO:0008354 | Germ cell migration |
| BP | GO:0006122 | Mitochondrial electron transport, ubiquinol to cytochrome c |
| BP | GO:0040003 | Chitin-based cuticle development |
| BP | GO:0055085 | Transmembrane transport |
| BP | GO:0055114 | Oxidation-reduction process* |
| BP | GO:0048142 | Germarium-derived cystoblast division |
| BP | GO:0007280 | Pole cell migration |

**Table 2.**
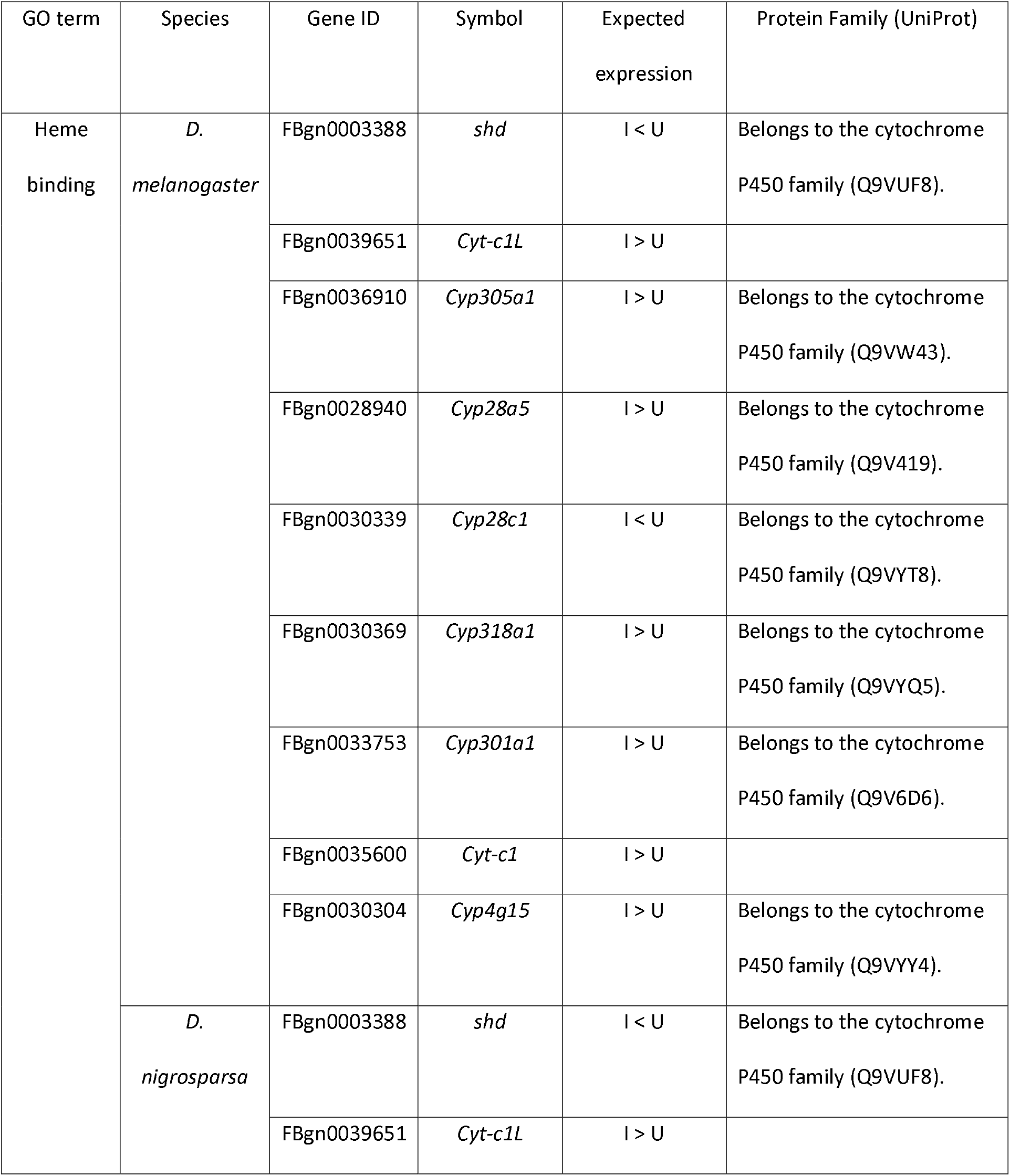
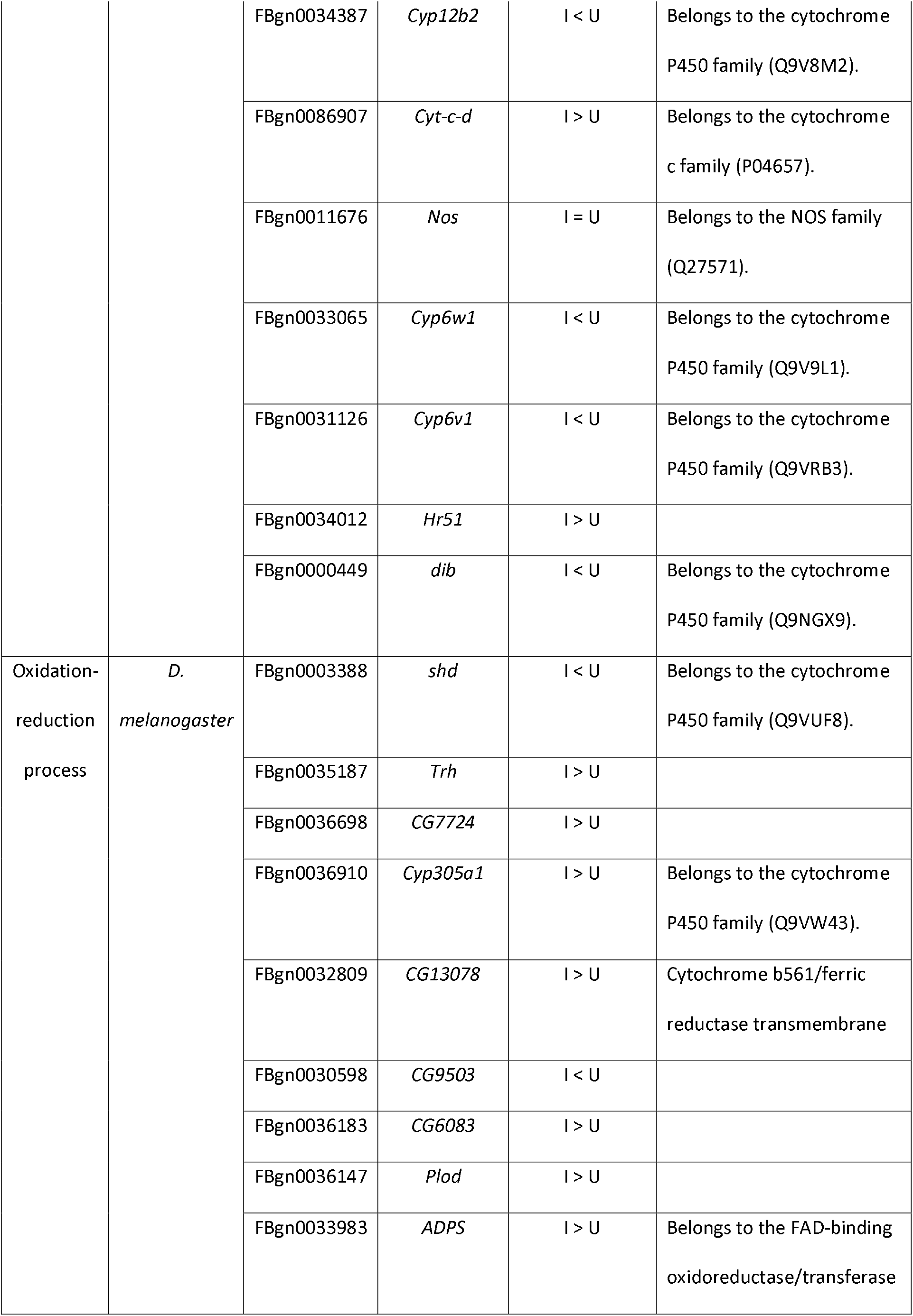
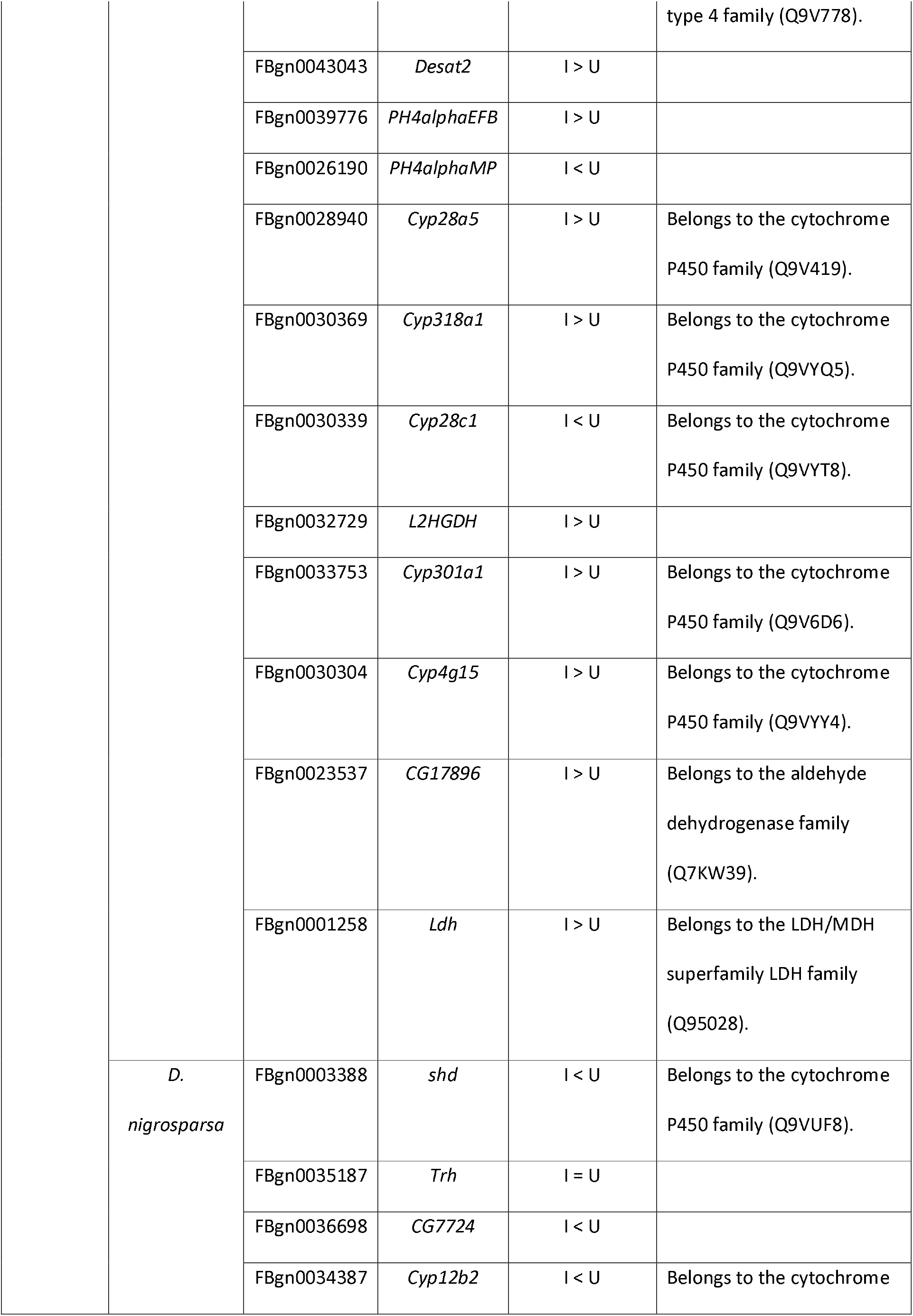
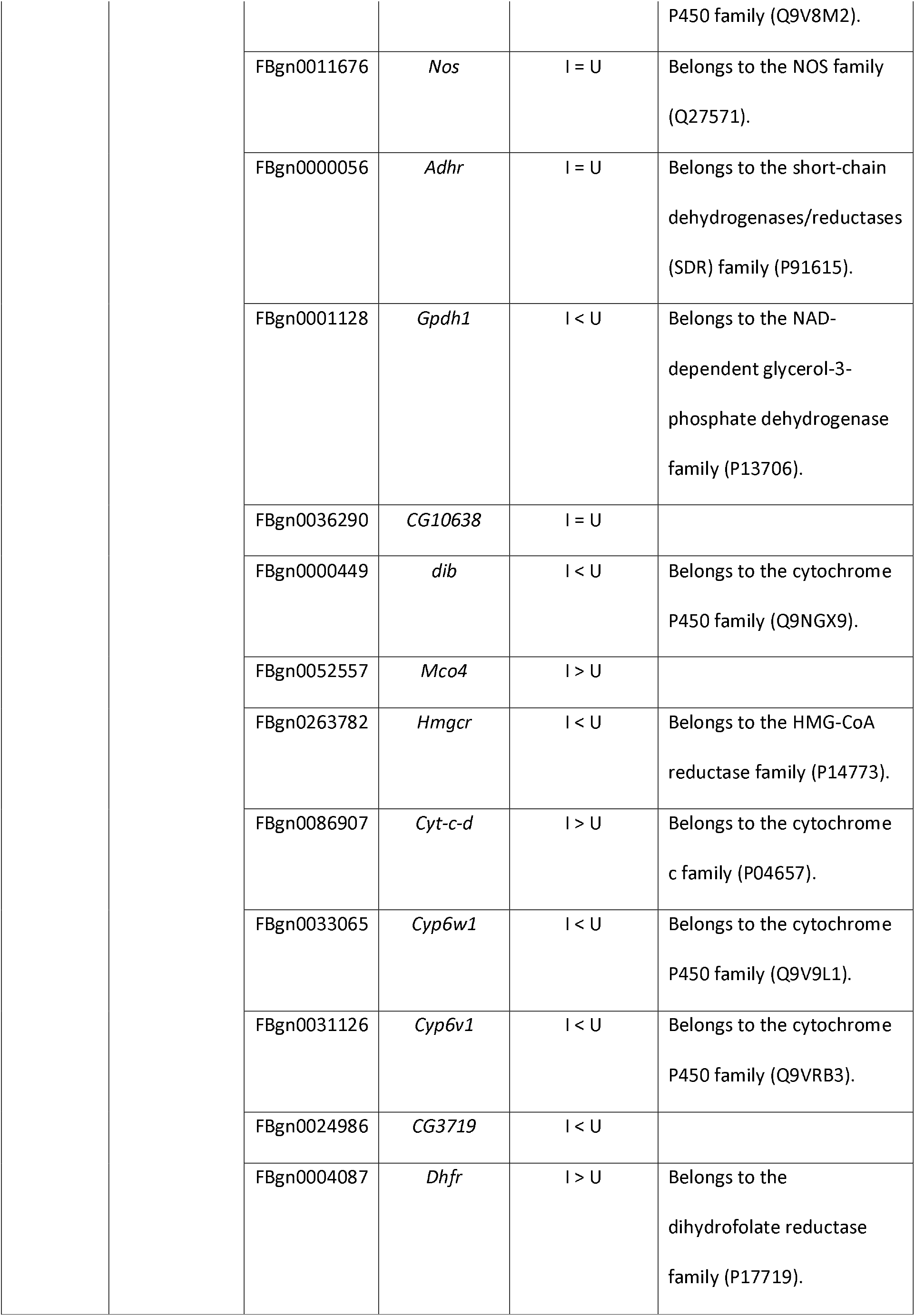
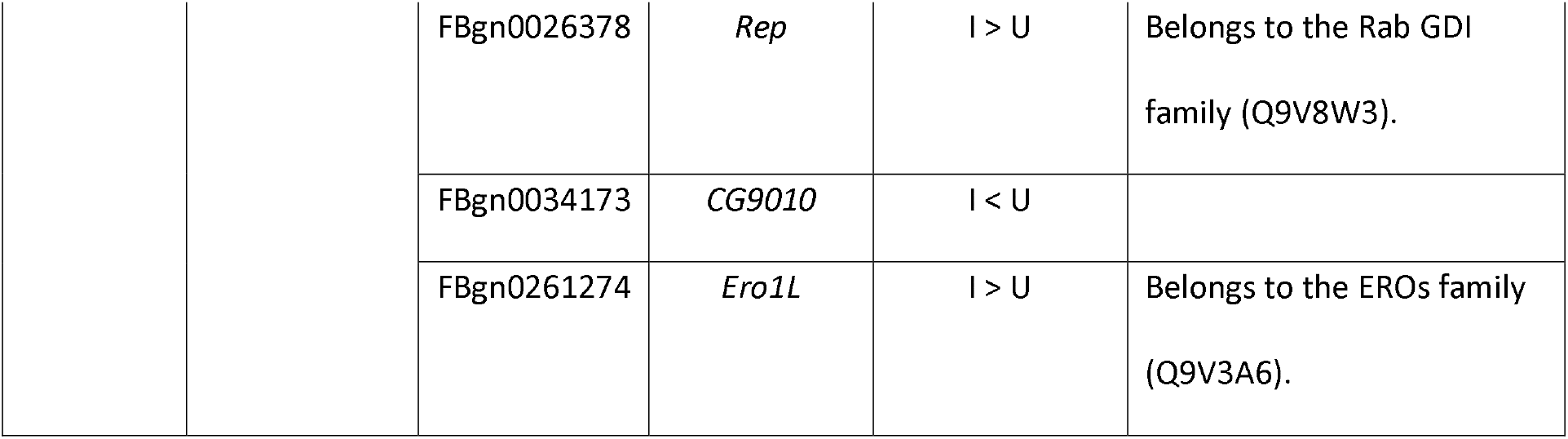
Expected gene expression between uninfected (U) and *Wolbachia*-infected (I), protein family, and molecular function of genes belonging to two shared gene ontology (GO) terms, heme binding (molecular function) and oxidation-reduction process (biological process), in *Drosophila melanogaster* and in *D. nigrosparsa*.

Three genes were shared between *D. melanogaster* and *D. nigrosparsa*, belonging to the biological process oxidation-reduction process, namely *shd*, *Trh*, and *CG7724*. These genes were likely to be expressed differentially between species (Table 2). Most genes were likely to be expressed more highly in infected *D. melanogaster* compared with uninfected lines of the same species.

## Discussion

High-throughput RNA sequencing offers the opportunity to detect and identify candidate genes that respond to *Wolbachia* infection. We found that gene expression was largely species-specific in *Drosophila melanogaster* and *D. nigrosparsa*, both when uninfected and infected with the *w*Mel strain of *Wolbachia*. In detail, however, we found a few differences in infection-related differential gene expression.

Independently of hosts’ particular phylogenetic position and ecology, the same *Wolbachia* strain can have different effects on different genetic backgrounds within and across host species (Ranz *et al*., 2003; Herbert and McGraw, 2018). A recent study in three black fly species in the genus *Simulium* found differential *Wolbachia* prevalence among species, suggesting host-specific interactions (Woodford *et al*., 2018). However, the mechanisms behind these interactions are not clear. Additionally, failure to transinfect *Wolbachia* strains from their native hosts to other species has been shown in many species, for example, between related species of parasitic wasps in the genus *Trichogramma* (Huigens *et al*., 2004) and from *D. melanogaster* to *D. nigrosparsa* (Detcharoen *et al*., 2020).

In our data, we found variation among pools of individuals within replicate lines and among replicate lines within species. With regard to the variation among pools of individuals within replicate lines, gene expression can be influenced by several factors such as the environment and individual variation (Wittkopp, 2007). For example, around 23 percent of the genes in *D. melanogaster* are expressed differently at the individual level, which could be due to individual variation in size or weight (Lin *et al*., 2016). In our study, we cannot evaluate an impact of individual variation as we used a pool of five individual flies per replicate, but, in any case, we would perhaps have seen less variation among pools if we had pooled more than five individuals per replicate.

Variation among replicate lines might result from several processes such as genetic drift and inbreeding, (Kristensen *et al*., 2006; Dunning *et al*., 2014). The lines of *D. melanogaster* and *D. nigrosparsa* were kept at a census size of 200 and 100 flies per line, respectively. The *D. melanogaster* lines used in our study were separated from outbred populations about two years before the experiment, whereas each infected *D. nigrosparsa* line was derived from a single transinfected female and kept separately for about two years. We suggest that the variation in gene expression among replicate lines may be due to a combination of both drift and inbreeding. In detail, the replicate lines in *D. nigrosparsa* were more separated than in *D. melanogaster* (Figure S2: R_line_ = 0.33 and 0.14 in values *D. nigrosparsa* and *D. melanogaster*, respectively), which would be in line with a smaller effective population size due to the stronger bottleneck in *D. nigrosparsa* and thus stronger effects of drift.

Apart from species-specific gene expression and variation among pools of individuals and replicate lines, we found that two GO-terms were affected in the two species but that the hosts reacted differently to the infection. Many genes involved in the oxidation-reduction process function as oxidoreductase activators (Table 2). Oxidoreductases produce reactive oxygen species (ROS) as a by-product in the oxidation-reduction process. However, these enzymes can also help in redox homeostasis of the host. *Wolbachia* can help their hosts in the redox homeostasis by producing their own oxidoreductase (Kurz et al, 2009), which, in turn, reduces the expression of the hosts’ genes. We found that multiple genes belonging to the oxidation-reduction process were up-regulated in the infected *D. melanogaster* but down-regulated in the infected *D. nigrosparsa* (Table 2). The low expression of these genes in infected *D. nigrosparsa* might indicate effects on this species in terms of protection against ROS.

ROS, in addition to being a by-product of molecular respiration, can be produced by heme in its free state (Oliveira *et al*., 1999; Paiva-Silva *et al*., 2006). Binding and degrading the ROS-induced heme are used in insects to regulate heme homeostasis (Paiva-Silva *et al*., 2006). Iron is released when the heme is degraded, but an excess of iron is harmful to organisms. *Wolbachia* reduce the iron concentration in host cells by producing proteins that help with iron storage, which as a result changes iron-related gene expression in hosts (Kremer *et al*., 2009, 2012). Here, genes involved in heme binding were expressed differently in the two Drosophila species (Table 2). Infected *D. melanogaster* expressed more of these genes than infected *D. nigrosparsa*. This finding on heme-binding related genes implies that the infection might benefit *D. nigrosparsa* in the heme-related process, similarly as in the oxidation-reduction process.

In summary, we conclude that the infection by the same *Wolbachia* strain induced a differential response in *D. melanogaster* and in *D. nigrosparsa*. Different gene expression patterns between species might be due to host specificity of the *Wolbachia* strain *w*Mel. We found two shared GO-terms, heme binding and oxidation-reduction process, shared between the two species. Down-regulated expression of genes in these two GO-terms in the infected-*D. nigrosparsa* may indicate small positive effects of *Wolbachia* on *D. nigrosparsa*. A long-term study should be done to observe changes in differential gene expression and to validate whether *D. nigrosparsa* would benefit from the infection. In addition, our results here were derived from whole-body extraction, but expression in specific tissues such as reproductive tissue should be tested in further studies for a better understanding of *Wolbachia*-induced gene expression.

## Acknowledgments

We thank Luis Teixera for providing *D. melanogaster* lines; Yuk-Sang Chan for performing microinjections; Philipp Andesner for helping with RNA extraction and quantification; Anja Ekblad for fly maintenance; MD was supported by the University of Innsbruck.

## Supplementary Data

**Figure S1.**
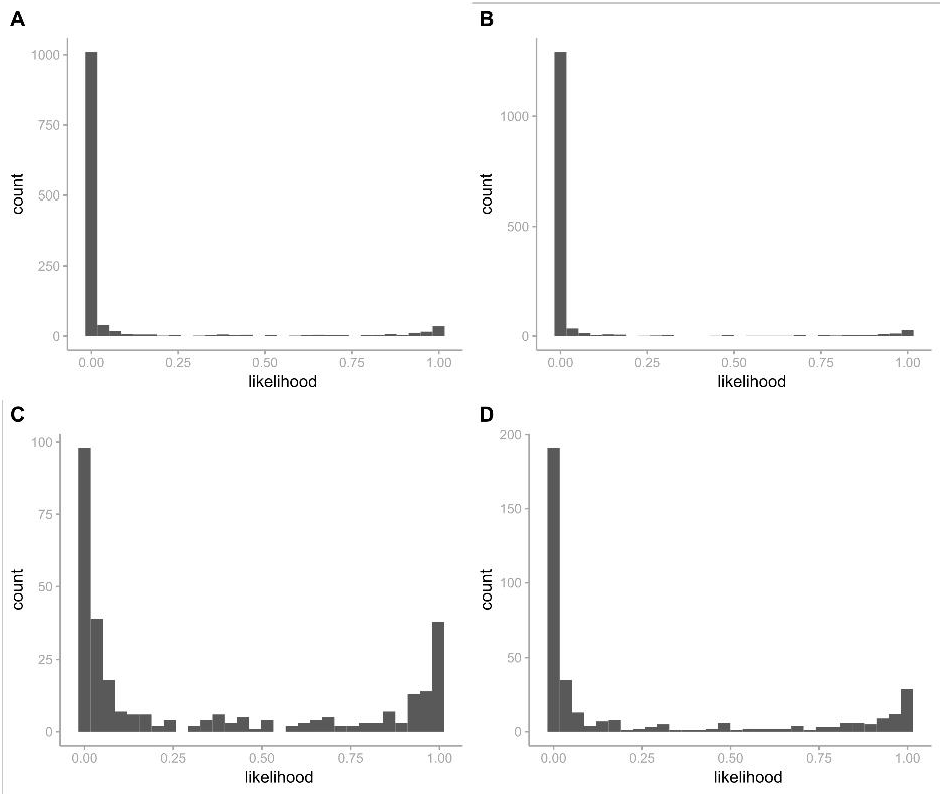

**Figure S2.**
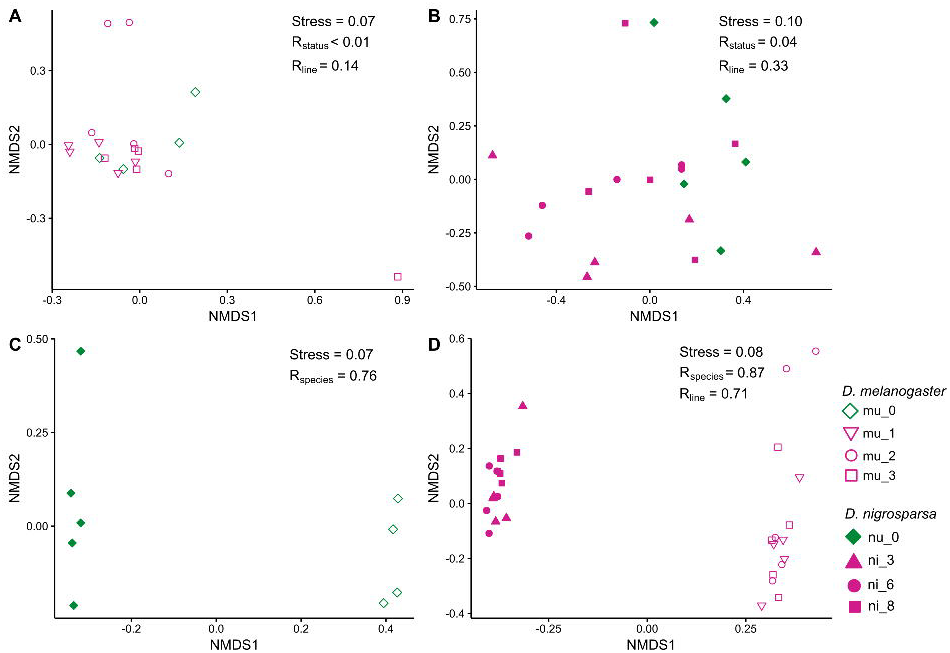

**Table S1.**
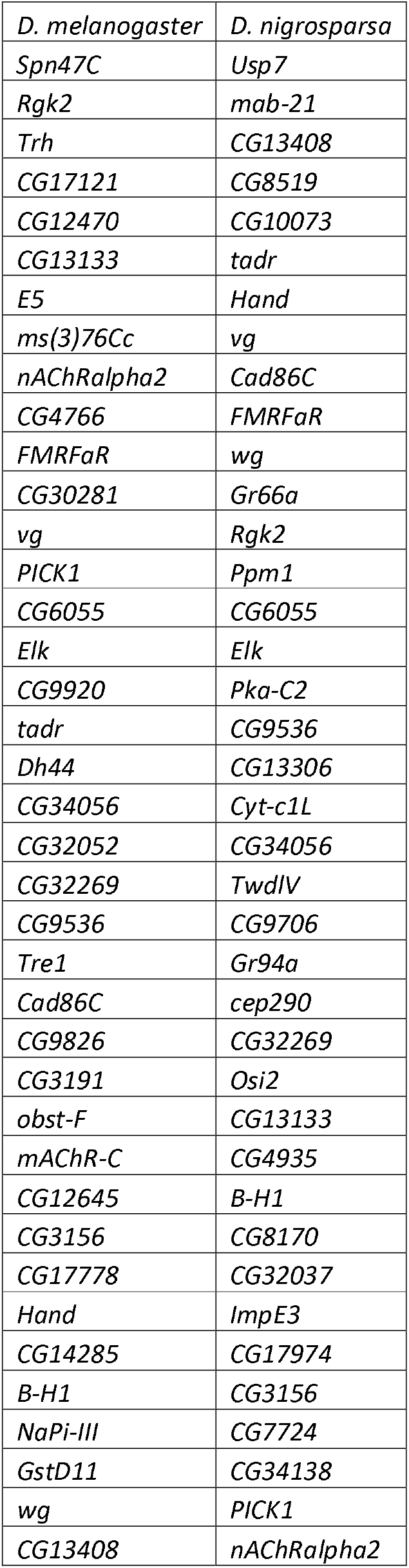
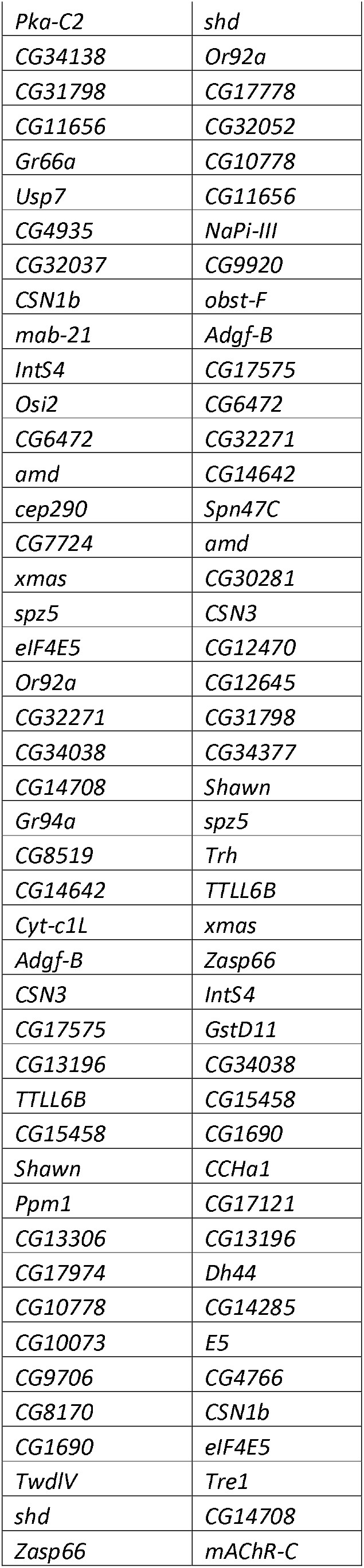
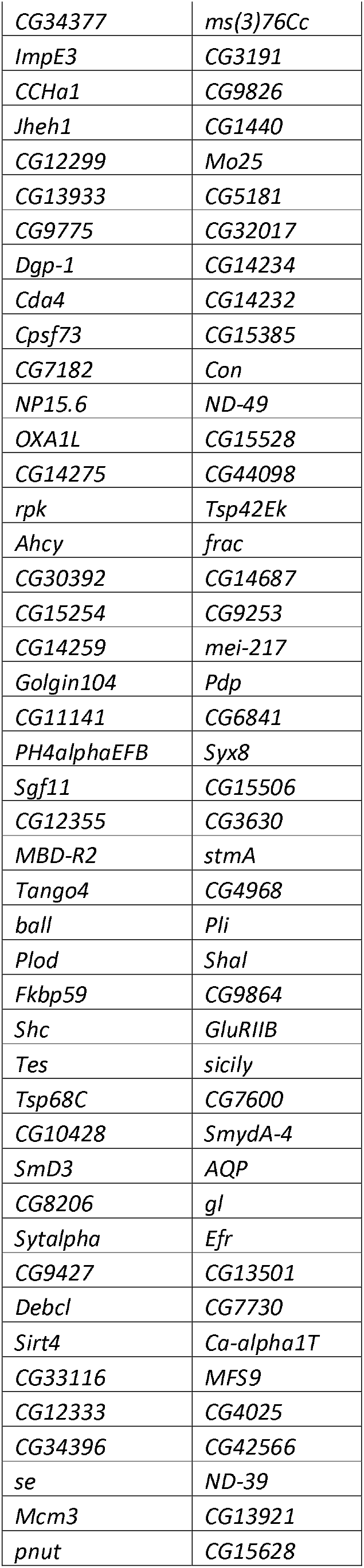
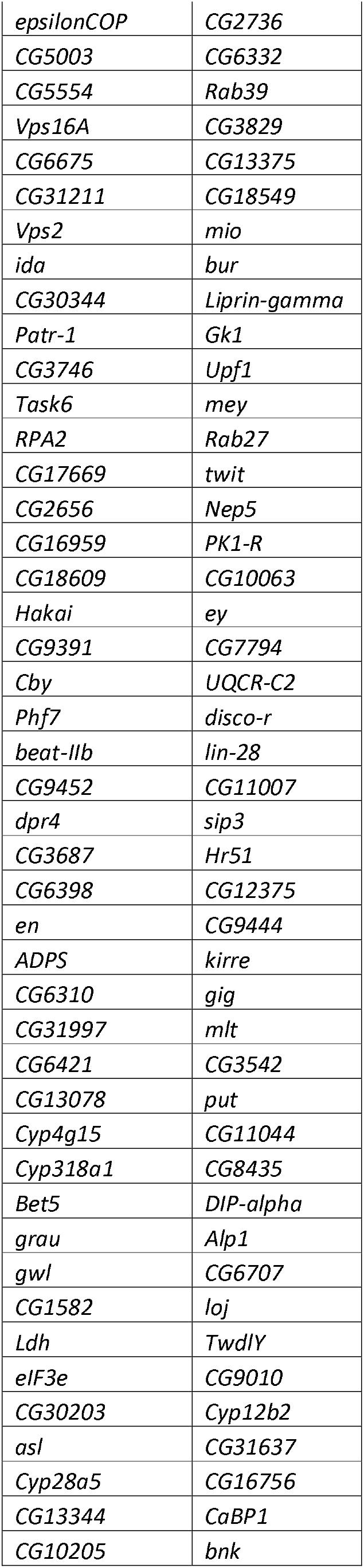
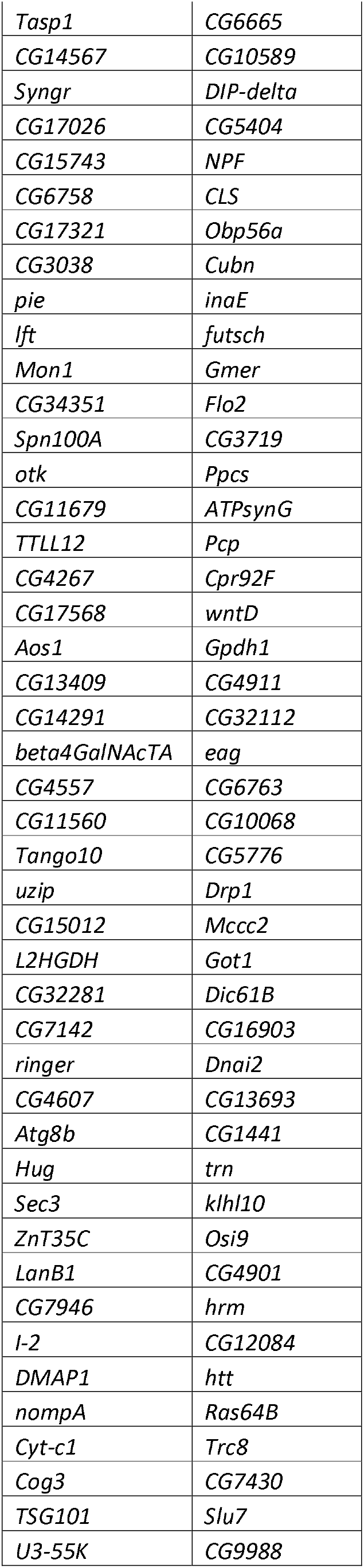
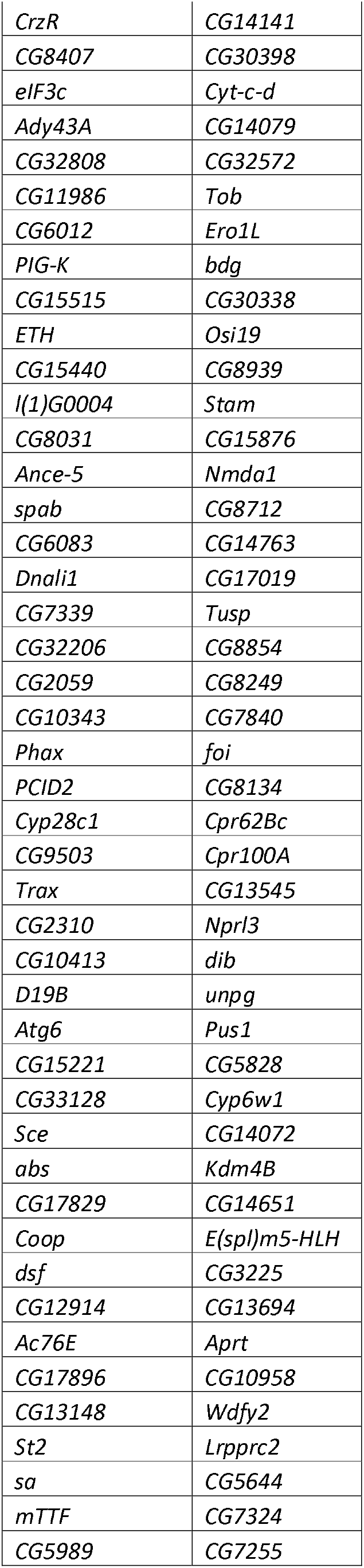
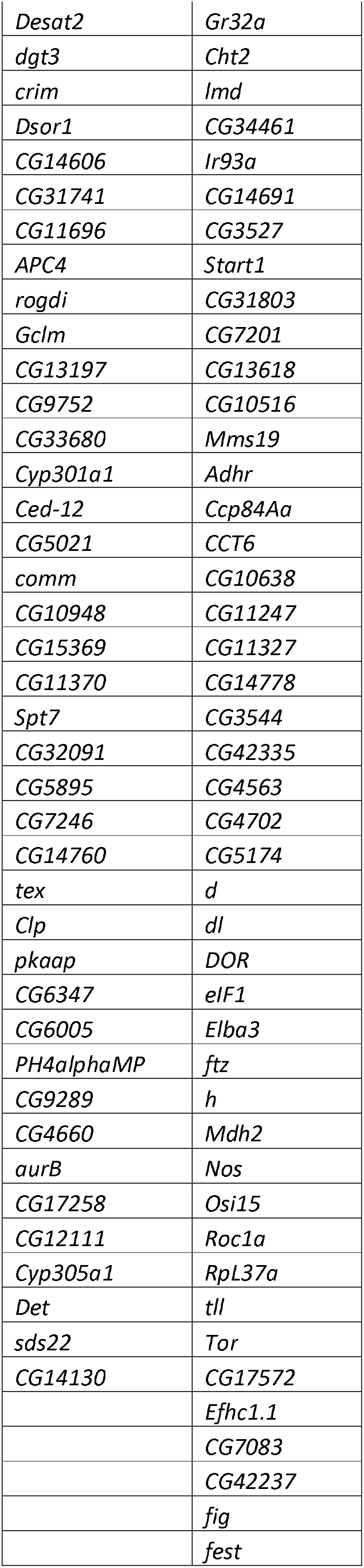
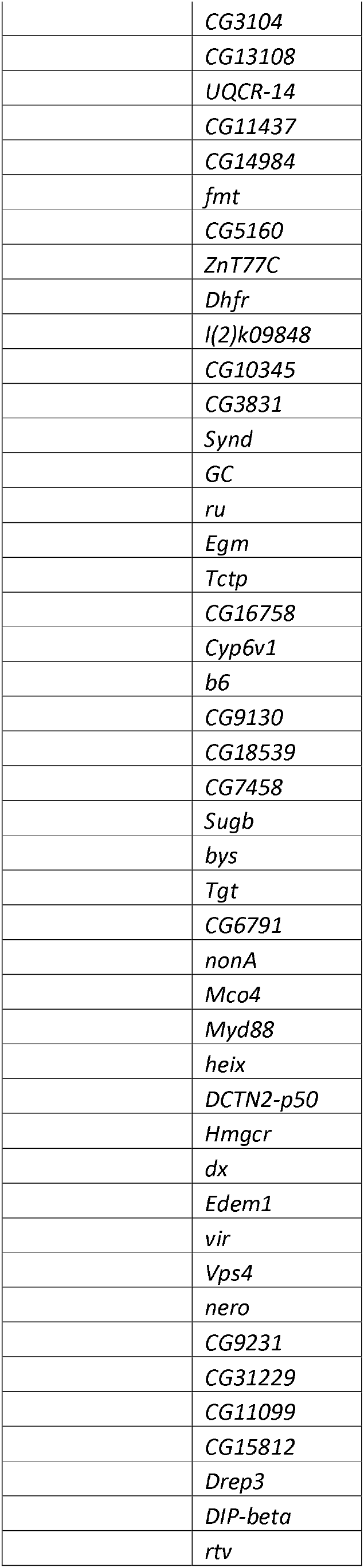
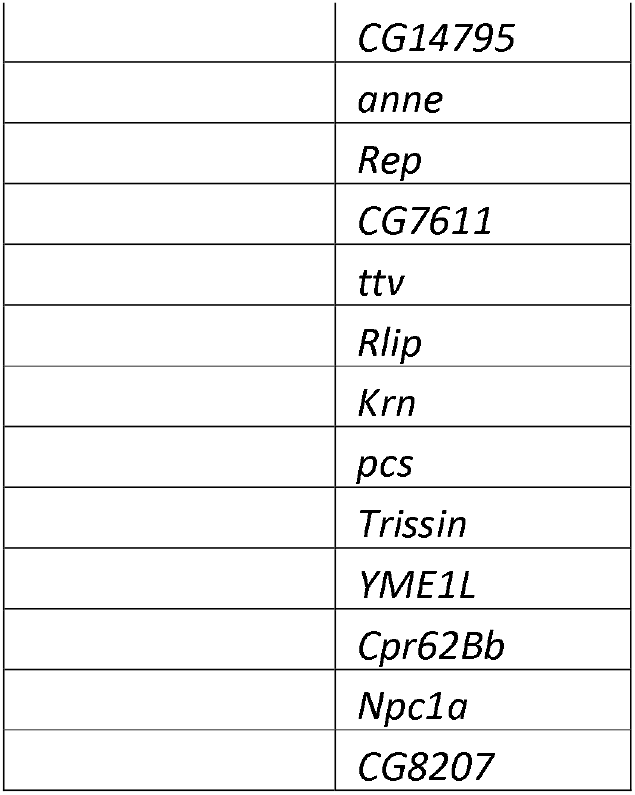
List of genes in the fourth quartile of *Drosophila melanogaster* and *D. nigrosparsa*.

